# Rapid terabase-scale simulation of realistic metagenomes for experimental design and pathogen detection with RandomReadsMG

**DOI:** 10.1101/2025.07.18.665570

**Authors:** Brian Bushnell, Frederik Schulz, Juan C. Villada

## Abstract

Accurate simulation of metagenomic sequencing data is determinant for benchmarking new algorithms, guiding experimental design and generation of synthetic data to develop and train models for AI/ML. However, existing simulators often struggle to reproduce the uneven coverage depth patterns seen in real microbial communities, can be difficult to install, or incur long runtimes. Here we present `RandomReadsMG`, a fast and scalable read simulator designed to generate synthetic metagenomes with realistic, user-defined depth distributions from hundreds of genomes in a single command. The tool supports multiple abundance models and intra-genome depth variability to better mimic real data. Benchmarks show `RandomReadsMG` runs orders of magnitude faster than comparable software while maintaining constant memory usage for input datasets of unbounded size. We demonstrate its utility by determining pathogen detection thresholds in a complex microbiome, showcasing its value for optimizing experimental design and creating robust training data sets for bioinformatics and AI/ML. `RandomReadsMG` is open-source software, distributed as part of the BBTools suite. The full software package is available for download at https://sourceforge.net/projects/bbmap/. For containerized deployment, a Docker image is also available from Docker Hub at https://hub.docker.com/r/bryce911/bbtools.

## 2. Introduction

The analysis of metagenomic data is fundamental to understanding microbial communities (Riesenfeld et al., 2004), but it presents significant computational challenges (Zhou et al., 2015), including massive dataset sizes and the lack of a known ground truth for real-world samples. The absence of such data sets hampers the development and validation of new bioinformatics tools. For critical applications like metagenomic binning, simply shredding reference genomes is an inadequate method for creating benchmark data, as it fails to reproduce the variable read coverage and depth patterns that are essential for defining genome boundaries. The generation of realistic, synthetic datasets with known composition is therefore determinant for benchmarking new algorithms, guiding experimental design, and training artificial intelligence (AI)/ machine learning (ML) models (Gonzales et al., 2023).

A variety of read simulators have been developed to address this need, including tools like `CAMISIM`(Fritz et al., 2019), `MeSS` (Chaabane et al., 2024), `InSilicoSeq` (Gourlé et al., 2019), `NEAT` (Stephens et al., 2016), `wgsim` (Li et al., 2009), `Mason` (Holtgrewe, 2010), `ART` (Huang et al., 2012), `BEAR` (Johnson et al., 2014), `FASTQSim` (Shcherbina, 2014), and `MetaSim`(Richter et al., 2008). However, these are often designed with a focus on single-genome sequencing to model instrument-specific error profiles for applications like variant detection, and typically assume uniform read depth distribution. Adapting such tools for metagenomics is cumbersome, requiring users to execute hundreds of separate commands with custom parameters to simulate a community with diverse abundances. While metagenome-specific simulators such as `MetaSim`, `InSilicoSeq`, and `MeSS`exist, they can be limited by long runtimes, complex dependencies, or may not easily model the non-uniform intra-genome read depth distributions seen in real data. Key features for metagenomic simulation, such as custom headers to retain ground-truth taxonomic origin or the ability to easily spike in organisms at specific low abundances, are often not primary design considerations.

Here we present `RandomReadsMG`, a fast and scalable read simulator designed specifically for generating complex metagenomic datasets. `RandomReadsMG`streamlines the simulation process, enabling the creation of a library from thousands of genomes with realistic abundance distributions in a single command. It supports multiple inter-genome coverage models to mimic the long-tailed species distributions common in nature, as well as configurable intra-genome coverage variability to better emulate real sequencing data. Recognizing that complex error modeling can be a bottleneck for generating the large volumes of data needed for modern AI/ML applications, `RandomReadsMG` makes these features optional, prioritizing high-speed data generation. As part of the widely used, dependency-free BBTools suite, `RandomReadsMG` provides an accessible, powerful tool for the bioinformatics community.

## 3. Results

### 3.1. RandomReadsMG provides a flexible workflow for realistic metagenome simulation

`RandomReadsMG` is engineered to generate synthetic metagenomic sequencing data with a high degree of fidelity and user control (Figure 1A). The tool’s comprehensive workflow, summarized in Figure 1A, guides the user from the input of reference genomes in FASTA format to the output of simulated reads in FASTQ format. The simulation process is governed by a modular set of parameters that allow for precise control over key aspects of the synthetic dataset. Users can define genome-level coverage using several abundance distributions (e.g., LINEAR, EXPONENTIAL) or assign custom depths to individual genomes, a feature useful in cases like for example spiking in organisms of interest like pathogens. The framework supports major sequencing platforms, including Illumina, Oxford Nanopore (ONT), and PacBio HiFi, by modeling their distinct read length profiles, insert size distributions, and error patterns.

**Figure 1.**
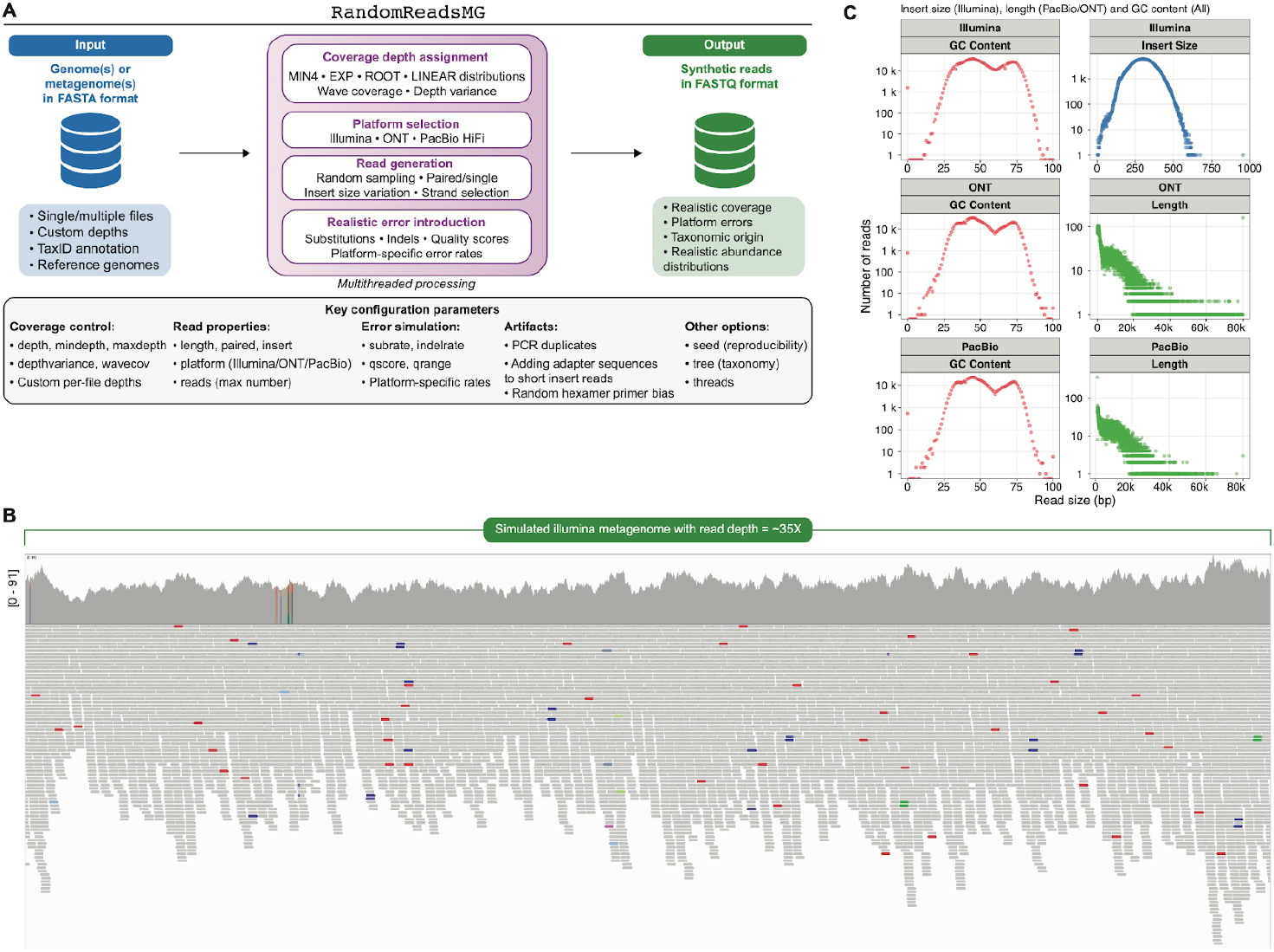
Overview of the RandomReadsMG simulation workflow and output characteristics. **(A)** Schematic diagram of the `RandomReadsMG` workflow, detailing the path from input FASTA files to output FASTQ reads. Key configurable modules include coverage modeling, platform selection, and the simulation of errors and artifacts. **(B)** A representative coverage plot from a simulated Illumina metagenome. The figure shows the variable, wave-like read depth generated along a contig. Visualization was done with the Integrative Genomics Viewer (IGV) (Robinson et al., 2011). **(C)** Simulated GC, insert size (Illumina), and length (Pacbio/ONT) distributions to demonstrate the simultaneous modeling of both compositional and structural read properties modeled by `RandomReadsMG`.

A key feature of `RandomReadsMG` is its ability to emulate the non-uniform read depth distribution frequently observed in real sequencing data. The tool can generate variable read depth within a single genome using a sine-wave-sum model, which prevents the artificially uniform coverage produced by simpler simulators. As presented in Figure 1B, `RandomReadsMG` provides a realistic, unpredictable coverage distribution across a genome in a simulated Illumina metagenome dataset. This intra-genome variability is essential for developing and benchmarking algorithms, such as metagenome binners, that rely on coverage patterns.

Furthermore, the fidelity of the simulation of `RandomReadsMG` extends to multiple platform-specific read characteristics. As shown in Figure 1C, `RandomReadsMG` models not only the characteristic read length and insert size profiles for Illumina, ONT, and PacBio platforms, with a GC content distribution reflective of the community and sequence lengths. The simulator accurately reproduces the tight insert size distribution of Illumina libraries and the broad, heavy-tailed read length distributions typical of ONT and PacBio sequencing. Additionally modeled characteristics such as random hexamer priming bias and PCR duplicates allow for the creation of complex, realistic, and reproducible synthetic datasets for a wide range of bioinformatics applications.

### 3.2. RandomReadsMG exhibits superior performance and scalability

To assess the computational performance of `RandomReadsMG`, we benchmarked its data generation speed and efficiency against other relatively modern read simulators, including `InSilicoSeq`, `wgsim`, and `RandomReads`. The benchmark was performed by simulating 2 million 2×150bp read pairs from a single-contig, 4.5Mbp bacterial genome (600 Mbp of data), with an additional test for `RandomReadsMG` using 192 genomes (15.053 Gbp of data) to evaluate its multi-genome performance. As presented in Figure 2A, `RandomReadsMG` demonstrates substantially higher throughput and efficiency. In single-genome mode, it generated data at 243.4 Mbp/s, a rate roughly 4.5 times faster than the next fastest tool, `wgsim` (54.5 Mbp/s). This performance advantage is even more pronounced in its native multi-genome mode, where its multithreaded-by-file design achieves a speed of 402.8 Mbp/s. A similar trend was observed for CPU efficiency, where `RandomReadsMG` was more than twice as efficient as `wgsim` and orders of magnitude more efficient than `InSilicoSeq`. See Supplemental Table S1 for more information. CAMISIM (Fritz et al., 2019) was deliberately not evaluated as it functions primarily as a wrapper for existing tools such as `wgsim` rather than an independent simulator, limiting its utility for comparative analysis.

**Figure 2.**
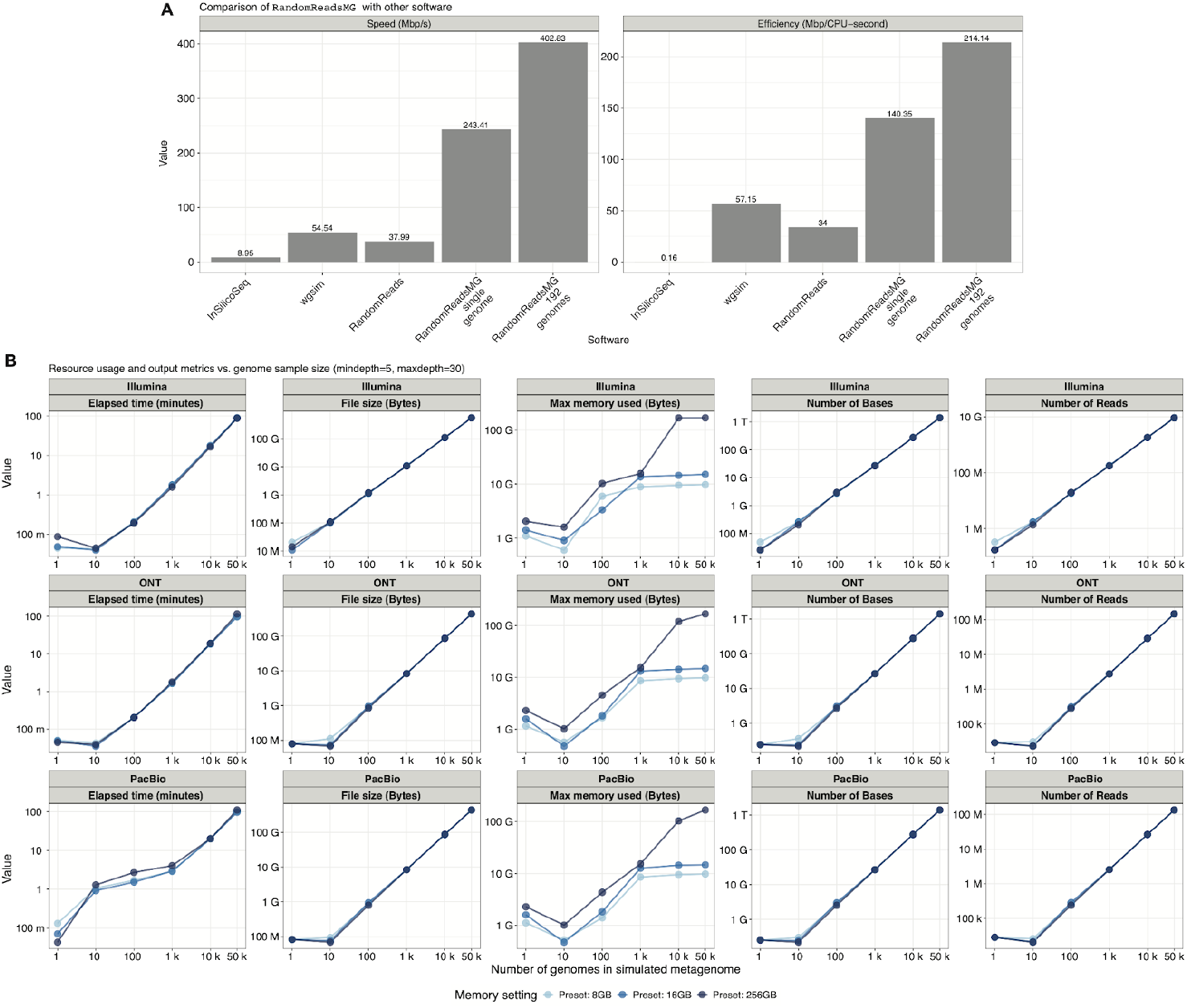
Performance benchmarks and resource scalability of RandomReadsMG. **(A)** Comparison of `RandomReadsMG` with other read simulators (`InSilicoSeq`, `wgsim`, `RandomReads`). The left panel shows data generation speed (Mbp/s) and the right panel shows CPU efficiency (Mbp/CPU-second). `RandomReadsMG` was tested in both single-genome and multi-genome (192 genomes) modes. **(B)** Resource usage and output metrics as a function of the number of input genomes (from 1 to 50,000) for Illumina, ONT, and PacBio simulations. Plots show elapsed time, output file size, maximum memory usage, and the total number of bases and reads generated. Lines correspond to different user-defined memory presets, demonstrating memory-constrained operation.

The scalability and resource consumption of `RandomReadsMG` were further evaluated by simulating datasets of increasing complexity, from 1 to 50,000 input genomes, for Illumina, ONT, and PacBio platforms (Figure 2B). This analysis shows that key metrics, including elapsed time, output file size, and the total number of reads and bases generated, scale in a predictable, linear fashion with the number of genomes included in the simulation. A critical aspect of `RandomReadsMG` design is memory management, as memory requirements can become a bottleneck in large-scale simulations. The results demonstrate that maximum memory usage is effectively controlled by user-defined presets. While memory consumption increases with the number of input files, it adheres to the specified limits, allowing the tool to operate robustly on systems with varying memory capacities. This predictable scaling and efficient resource management make `RandomReadsMG` well-suited for generating the massive, complex datasets required for developing and training modern bioinformatics tools, even in laptops with low hardware resources.

To further dissect the tool’s resource handling, we analyzed its memory efficiency relative to the user-specified allocation across simulations of increasing complexity (Supplemental Figure 1). This analysis reveals that for simulations involving a small number of genomes, memory efficiency is modest, as the tool’s memory footprint is significantly smaller than the allocated amount. However, as the number of input genomes increases into the thousands, the actual memory usage rises to meet the preset limit, without negatively impacting speed. This demonstrates that while `RandomReadsMG` operates well within its allocated memory ceiling, it also effectively utilizes the available resources when simulating large and complex metagenomes. The actual memory consumed scales predictably with the number of genomes until the user-defined cap is reached, at which point it plateaus, ensuring stable performance without memory over-consumption.

**Supplementary Figure 1.**
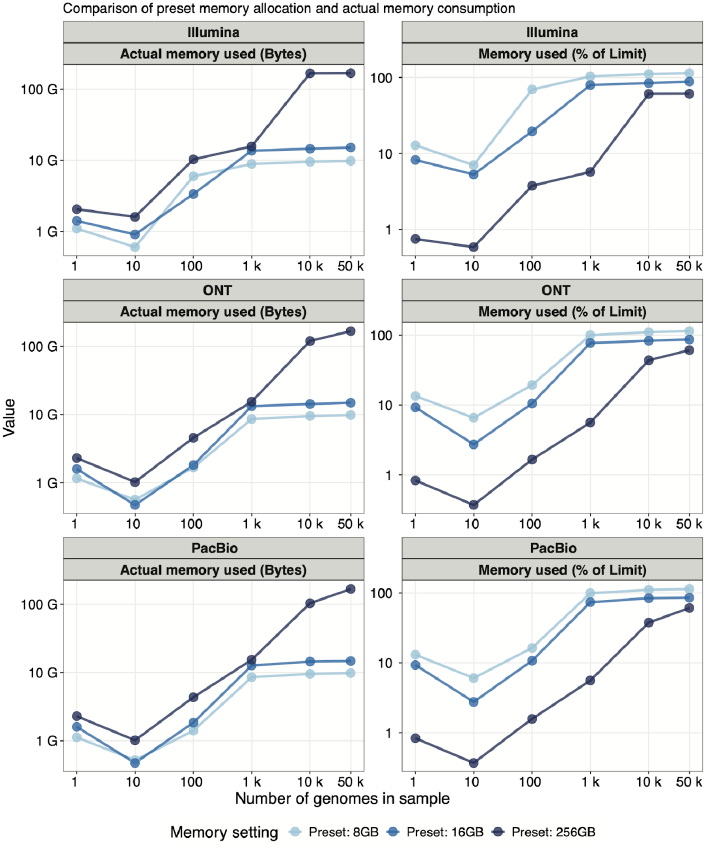
Comparison of preset memory allocation and actual memory consumption. Actual memory used in bytes (left column) and memory efficiency (right column) as a function of the number of genomes in the simulated sample for Illumina, ONT, and PacBio platforms. Lines correspond to runs with different Java memory limits(-Xmx8G, -Xmx16G, and -Xmx256G). The plots show that memory usage scales with the number of input genomes until the user-defined limit is reached.

### 3.3. RandomReadsMG for determining pathogen detection thresholds in drinking water microbiomes

To demonstrate the practical utility of `RandomReadsMG` in guiding experimental design for biosurveillance, we performed an in silico experiment to determine the sequencing depth required to detect and recover genomes of common waterborne pathogens from a realistic, complex microbial background. For this purpose, we first constructed a representative baseline drinking water (DW) microbiome to serve as a challenging and realistic community for our simulations (Figure 3A). This community was built by combining 77 high-quality metagenome-assembled genomes (MAGs) from a published drinking water study (Sevillano et al., 2021) with 26 representative species genomes from NCBI, resulting in a dereplicated set of 103 unique genomes (Supplemental Table S2). To ensure the simulated background reflected a natural abundance profile rather than an arbitrary uniform distribution, we mapped reads from 33 real DW metagenomes (Supplemental Table S3) to this genome set to establish a baseline coverage for each member. The resulting baseline community is taxonomically diverse, dominated by members of Pseudomonadota, and varied in its genomic characteristics, providing a suitable background for pathogen detection experiments (Figure 3B).

**Figure 3.**
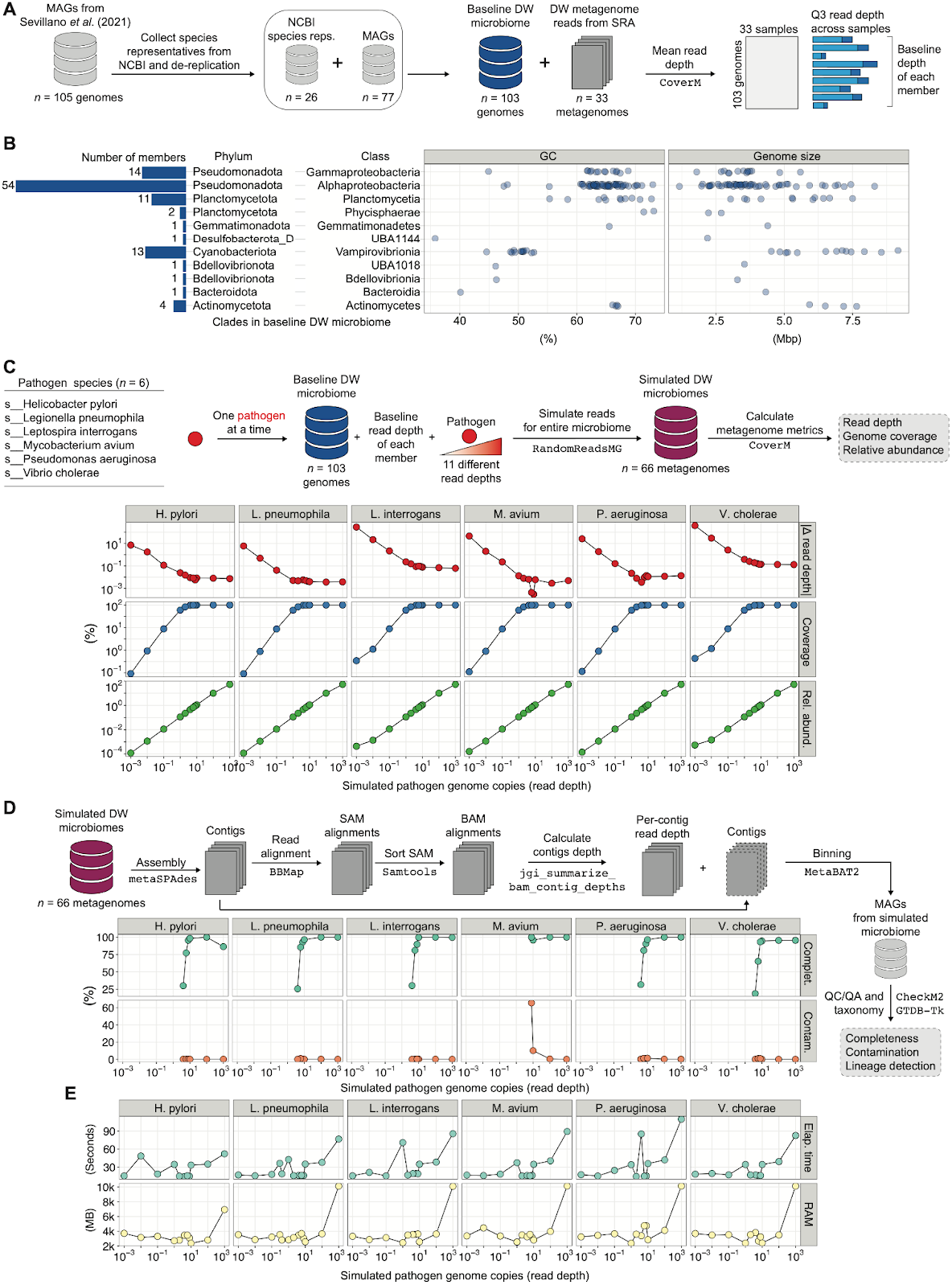
Simulation of pathogen detection in a drinking water microbiome. **(A)** Workflow for the construction of a baseline drinking water (DW) microbiome, composed of 103 genomes with realistic abundance profiles derived from real metagenomic data. **(B)** Taxonomic and genomic characteristics of the 103 members of the baseline DW microbiome. **(C)** Validation of the simulation parameters. Six different pathogens were spiked into the baseline community at 11 different read depths. The resulting read depth, genome coverage, and relative abundance are shown to correlate directly with the simulated input depth. **(D)** Assessment of pathogen genome recovery. Each simulated metagenome was processed through an assembly and binning pipeline (top schematic). The completeness and contamination of the resulting pathogen MAG are plotted against the initial simulated pathogen depth (bottom plots). **(E)** Computational resources (elapsed time and RAM) required for the simulation of each metagenome.

Using this baseline DW community, we then simulated a series of metagenomes to test detection limits. We selected six bacterial species of public health concern (*Helicobacter pylori, Legionella pneumophila, Leptospira interrogans, Mycobacterium avium, Pseudomonas aeruginosa, Vibrio cholerae*) and, using `RandomReadsMG`, spiked each pathogen individually into the 103-member baseline community at 11 distinct read depths, ranging from extremely low (0.001x) to moderate (10x) coverage. This generated a total of 66 synthetic metagenomes, each representing a different detection scenario. We first validated the output of `RandomReadsMG` by measuring the resulting average read depth, genome coverage, and relative abundance of the spiked pathogen in each simulated dataset. As shown in Figure 3C, the empirical metrics for each pathogen scaled perfectly with the target simulation depth, confirming that `RandomReadsMG` accurately generates reads reflecting the desired fractional abundances, a prerequisite for conducting reliable sensitivity analyses.

Each of the 66 simulated metagenomes was then processed through a standard bioinformatics workflow, including metagenomic assembly, read mapping, and binning, to assess the feasibility of recovering the pathogen genome (Figure 3D, top panel). The quality of the resulting pathogen MAG from each simulation was evaluated for completeness and contamination. The results clearly demonstrate that the ability to recover a high-quality MAG is directly dependent on the simulated sequencing depth of the pathogen (Figure 3D). For most of the tested pathogens, a simulated depth of approximately 6x was necessary to reconstruct a MAG of >50% completeness and <5% contamination, with the exception of Mycobacterium avium which required approximately 100x; to reconstruct a MAG of >90% completeness and <1% contamination, depth of approximately 8x was necessary for *Helicobacter pylori, Legionella pneumophila, Pseudomonas aeruginosa*, and *Vibrio cholerae*, 10x for *Leptospira interrogans*, and 100x for *Mycobacterium avium*. Below this threshold, genome recovery was inconsistent, yielding highly fragmented assemblies and low-quality MAGs, if any. Hence, our results provide a quantitative estimate for the minimum sequencing coverage required for reliable pathogen genome reconstruction in a complex sample, showcasing how `RandomReadsMG` can be used to inform sequencing strategies and manage research budgets effectively.

Finally, we assessed the computational resources required by `RandomReadsMG` to perform the simulation of each dataset (Figure 3E). The total elapsed time and memory (RAM) usage for the complete simulations were recorded for each of the 66 metagenomes. The analysis remained computationally tractable across all simulations, with time and memory scaling predictably with the complexity of the dataset. Our application demonstrates the practicality of using `RandomReadsMG` to generate large and diverse sets of simulated data for systematically benchmarking bioinformatics pipelines or training machine learning models, as the subsequent analyses are computationally feasible.

## 4. Discussion

Here we have introduced `RandomReadsMG`, a high-performance metagenome read simulator designed to address the critical need for realistic, large-scale synthetic datasets. The limitations of existing tools (namely challenges in modeling realistic coverage distributions, cumbersome workflows for multi-genome simulation, and computational speed) create a bottleneck for the development and validation of bioinformatics software. `RandomReadsMG` overcomes these limitations through a metagenome-first design that prioritizes speed, scalability, and the faithful representation of community abundance structures, as demonstrated by our performance benchmarks (Figure 2) and pathogen detection use case (Figure 3).

A major application for `RandomReadsMG` is the creation of controlled, reproducible benchmarks for bioinformatics tools. The robust and objective assessment of software performance is a persistent challenge in computational biology, where outcomes can be highly sensitive to the nuances of input data (Meyer et al., 2022; Sczyrba et al., 2017). By enabling the precise simulation of key metagenomic features, such as fractional member (pathogen) abundance and non-uniform coverage depth, `RandomReadsMG` allows for the creation of standardized gold standard datasets. As demonstrated by community-wide benchmarking efforts like the Critical Assessment of Metagenome Interpretation (CAMI), such datasets are invaluable for ensuring fair and reproducible comparisons between different algorithms, from assemblers to binners to taxonomic profilers (Meyer et al., 2022; Sczyrba et al., 2017).

The ability to generate realistic data in silico has exciting implications for experimental design. Before committing significant resources to deep sequencing or extensive clinical sampling, researchers can use simulation to address fundamental questions about experimental power, such as the minimum sequencing depth required to detect a low-abundance pathogen or accurately profile a microbial community (Zaheer et al., 2018). As we demonstrated with our drinking water microbiome experiment, `RandomReadsMG` can be used to model these scenarios, allowing investigators to select cost-effective sequencing parameters that meet the sensitivity requirements of their study. This same capability is crucial for training robust Artificial Intelligence AI/ML models. The performance of such models is notoriously dependent on the quality and diversity of their training data, and models trained on overly simplistic or biased data often fail to generalize to real-world samples (Zou et al., 2019). `RandomReadsMG` facilitates the generation of hundreds of complex dataset replicates across a range of coverages, noise/error levels, and community structures, enabling the development of more accurate and generalizable AI/ML applications for metagenomics.

The ease of use and rapid performance of `RandomReadsMG` also serves to accelerate the entire cycle of scientific development and education. For algorithm developers, the ability to generate terabytes of test data in hours rather than weeks removes a significant bottleneck, allowing for faster iteration and validation of new methods. For educators, the tool provides a platform for hands-on exercises where students can interactively explore how sequencing artifacts and coverage biases affect downstream analysis, deepening their understanding of both sequencing technologies and bioinformatics principles (Attwood et al., 2019).

In conclusion, `RandomReadsMG` is a powerful and accessible tool that addresses an immediate and critical need in the bioinformatics community, empowering researchers to create better benchmarks, design smarter experiments, and ultimately accelerate the pace of discovery in microbiology while conserving both computational and physical resources.

## 5. Materials and methods

### 5.1. RandomReadsMG implementation

The version of `RandomReadsMG` used in this study is part of the BBTools suite v39.33. Like other BBTools, `RandomReadsMG` is written in pure Java with v1.8 compliance, and distributed with source code and precompiled class files, so it can run on any computer that supports Java. The program is multithreaded by input file; each can be read simultaneously, and have reads generated in an individual thread, which are then written to a shared output file (or two files for paired reads). This allows it to generate data at up to 402 Mbp/s, 10x faster than `RandomReads`. It supports FASTA or FASTQ input or output, optionally gzipped or bgzipped.

### 5.2. Coverage models

#### 5.2.1. Genome-level coverage

`RandomReadsMG` supports 4 inter-genome coverage models, with each genome assigned a random average coverage ranging from user-specified mindepth to maxdepth:

- MIN4: Minimum of four uniform random values
- EXP: Exponential distribution for long-tailed coverage
- ROOT: Square-root transformed distribution (depth=mindepth+(rand()^2)*range)
- LINEAR: Uniform random distribution
- Custom: Individual genomes may be assigned custom coverage

This allows customization depending on the profile of the data to simulate; the first three are all aimed at mimicking the low-depth-biased coverage of a typical metagenome. Additionally, individual input files can be assigned a custom depth to simulate a very low-level genome of interest, or a highly-dominant organism, or any desired coverage mix.

#### 5.2.2. Intra-genome base-level coverage

`RandomReadsMG` defaults to a simple model of intra-genome coverage with a linear gradient across contigs. This should be sufficient to simulate origin-of-replication bias or sequencing bias as it pertains to assembled contigs having nonuniform coverage for binning, clustering, or contamination-identification experiments.

A more sophisticated model is available via the “sinewave” flag. This varies the coverage realistically using a sum of multiple different-period sine waves, with the result looking something like the visualization of a sound file. The magnitude and number of sinewaves can be customized, and the specific waveform is randomly generated per sequence. A user-defined “minprob” parameter controls the minimum allowed relative coverage due to overlapping wave valleys, allowing coverage holes that prevent contiguous assemblies to be either prevented or promoted.

### 5.3. Sequencing platforms

`RandomReadsMG` supports 3 major read-sequencing platforms: Illumina, Oxford Nanopore (ONT), and Pacific Biosciences (PacBio) through preset models, via the “illumina”, “ont”, and “pacbio” flags, and can model similar platforms using custom settings for read length distributions and error rates. The default read lengths are: Illumina: 2×150 bp paired reads with a mean insert size of 300bp. ONT: Read lengths are taken from a mixed exponential-lognormal distribution with a heavy tail and mean length of 15 kbp. This is designed to capture both the predominant read distribution and the long tail containing the most valuable reads. PacBio HiFi: Read lengths taken from a log-normal distribution with mean length 15 kbp. These defaults can be overridden, including custom values for sigma, tailfactor, and min and max length.

### 5.4. Error simulation

Error simulations are specific for each platform. For short reads, the substitution errors (the overwhelmingly predominant type associated with Illumina sequencing) may be simulated by specifying a fixed or variable quality score, though variable quality scores lead to much larger gzipped output, and grant little advantage for most purposes. Specific positional quality profiles are not supported due to the high degree of variability in such profiles from instrument to instrument and even run to run, along with the fact that real sequence quality scores (unlike synthetic ones) tend to be extremely inaccurate anyway. The default quality score is 25 with 0 variance. For long reads, the PacBio and Nanopore modes include a mix of substitution, insertion, and deletion errors, with the default settings for each platform reflecting observed rates (Hifi/CCS rates, for PacBio). Homopolymers have an additional parameter that increases the chance of a homopolymer indel (deletion or insertion of the homopolymer base) as the stretch continues. All of these rates are customizable (see Table 1) to reflect future changes in chemistry and base-calling. Also, for both short and long reads, all error rates can be adjusted independently for any platform, but the homopolymer indel increase only affects long read mode.

**Table 1.** Default error rate table.

| Parameter | Illumina | ONT | PacBio |
| --- | --- | --- | --- |
| Quality score | 25 | 20 | 35 |
| Substitution rate | 0 | 0.0025 | 0.00015 |
| Insertion rate | 0 | 0.0055 | 0.000055 |
| Deletion rate | 0 | 0.0045 | 0.000045 |
| Homopolymer error | 0 | 0.02 | 0.000015 |

### 5.5. Other features

`RandomReadsMG` supports PCR duplicate simulation, with a specified duplication rate, for all platforms. PCR duplicates are explicitly labeled as duplicates (d_1 in the header with the original labeled d_0) and have the same genomic coordinates but independent sequencing error. User-definable adapter sequence may be automatically appended to fixed-length reads when the insert size is shorter than read length; after exhausting adapter sequence, the remaining length is filled with low-quality Poly-G, reflective of 2-dye Illumina platform behavior.

### 5.6. Comparison of RandomReadsMG performance

`wgsim`, `RandomReads`, and `RandomReadsMG` were each used to generate 2 million 2×150 bp read pairs from a single-contig, 4.5Mbp bacterial genome. Additionally, `RandomReadsMG` was used to generate random coverage from a directory containing 192 bacterial genomes. The command lines used, on a 2.5Ghz AMD EPYC 7502P 32-Core computer:

~~~
time wgsim -N2000000 -1150 -2150 -r0 ref.fa r1.fq r2.fq
time randomreads.sh reads=2m ref=ref.fa out=r#.fq len=150 paired
time randomreadsmg.sh reads=2m ref=ref.fa out=r#.fq len=150
time randomreadsmg.sh ∼/genomes out=r#.fq mindepth=10 maxdepth=40 time iss generate --genomes ref.fa --model hiseq --output r --n_reads
4761904 --cpus 32
~~~

Note: `RandomReadsMG` generated more data due to the way it operates in multi-genome mode. `InSilicoSeq` needed a different read count argument because it is limited to 126bp reads and the argument specifies read count unlike pair count in the case of the other tools.

`RandomReadsMG`‘s memory requirements scale with the number of threads (T) and input files used. For the active files, 1 byte of raw FASTA file on disk corresponds to 1 byte of RAM used (plus overhead), but only T files will be active at any time, so memory consumption is proportional to the T largest files in the input dataset. Furthermore, input files are streamed one sequence at a time, so with fragmented inputs, memory requirement is proportional to only T*(longest contig). By default T will equal the smaller of the number of files and logical processors but can be overridden with the “t=” flag to conserve memory.

Although not typically necessary, multiple threads can also be used to process a single file via the ‘singlefilethreads’ parameter, increasing speed when complex error rate and coverage settings are applied. In this case, again, the memory use scales linearly with the number of threads and the largest contig in the file.

### 5.7. RandomReadsMG usage

For ease of use and maximum reproducibility, RandomReadsMG is available as a Docker image. The version used in this manuscript (39.33) can be run, and the installation verified, with the following command, which will display the tool’s help menu::

~~~
docker run bryce911/bbtools:39.33 randomreadsmg.sh --help
~~~

`RandomReadsMG` has a simple command-line interface with an optional shellscript to help set Java parameters in Linux or MacOS environments.

Then, to generate reads from all of the FASTA files in directory “genomes” and write the output as 2×150 bp interleaved FASTQ to `reads.fq`:

~~~
randomreadsmg.sh genomes out=reads.fq
~~~

Or to specify a custom mixture of three files:

~~~
randomreadsmg.sh ecoli.fa=30 mruber.fa=0.01 phix.fa=100 out=reads.fq
~~~

Alternatively, for platforms such as Windows without native bash support, the equivalent Java command can be given:

~~~
java -ea -Xmx1g -cp C:/bbmap/current/bin.RandomReadsMG genomes
out=reads.fq
~~~

Further usage information and flag descriptions are available in `randomreadsmg.sh`, which will display usage information when run with no arguments, or opened in a text editor. The resultant reads are tagged with the TaxID of the originating file (if the file had a TaxID in the name). Thus, after assembling the reads, they can be aligned to the assembly and used to assign taxonomy; this is a simplified (but working) pipeline:

~~~
randomreadsmg.sh genomes out=reads.fq.gz
tadpole.sh in=reads.fq.gz out=contigs.fa k=93
bbmap.sh ref=contigs.fa in=reads.fq.gz out=mapped.sam.gz
renamebymapping.sh in=contigs.fa out=renamed.fa *.sam.gz
~~~

All BBTools are file extension-aware, so naming an output file `x.fa` or `x.fasta` will generate FASTA output, while `x.fq` or `x.fastq` would be FASTQ, and `.sam.gz`would be gzipped sam, etc. `RandomReadsMG` follows the standard `BBTools` conventions such as `in=stdin.fa` to stream FASTA input from standard in, and uses the same universal flags such as `zl` (compression level) and `fastawrap` to set the associated values.

### 5.8. Methods for pathogen detection sensitivity analysis

#### 5.8.1. Construction of a baseline drinking water microbiome (DWM)

To create a realistic and complex background for pathogen detection simulations, a baseline drinking water (DW) microbiome was constructed. The process began by sourcing microbial genomes from high-quality metagenome-assembled genomes (MAGs, obtained from https://doi.org/10.6084/m9.figshare.c.5414964.v2) from a previous study on drinking water metagenomics (Sevillano et al., 2021) and 26 representative species genomes from the National Center for Biotechnology Information (NCBI) (Kannan et al., 2023; Sayers et al., 2023). This collection of genomes (Supplemental Table S4) was then de-replicated to ensure a set of unique species representatives, resulting in a final baseline community of 103 distinct genomes. To model a natural abundance profile, we downloaded public sequencing data from 33 real drinking water metagenomes from the Sequence Read Archive (SRA). These reads were mapped against our 103-genome set, and the mean read depth for each genome across all 33 samples was calculated using `CoverM` (v0.7.0) (Aroney et al., 2025). The third quartile (Q3) of these mean depths was chosen as the baseline coverage for each respective member in our final synthetic community, ensuring a realistic, non-uniform abundance distribution.

#### 5.8.2. Simulation of pathogen-spiked metagenomes

The primary goal of this experiment was to determine the detection limits for specific pathogens within the constructed DW microbiome. We selected six bacterial species relevant to public health and water safety: *Helicobacter pylori, Legionella pneumophila, Leptospira interrogans, Mycobacterium avium, Pseudomonas aeruginosa*, and *Vibrio cholerae*. Using `RandomReadsMG`, we generated a series of synthetic metagenomes. In each simulation, the 103-member baseline DW community was included at its pre-calculated baseline coverage. Then, one pathogen at a time was spiked into this community at 11 different target read depths, ranging from 0.001x to 10x. This process was repeated for all six pathogens, resulting in a total of 66 distinct simulated metagenomes for downstream analysis. To confirm the fidelity of the simulations, the actual read depth, genome coverage, and relative abundance of the spiked pathogen in each output file were calculated using `CoverM` and compared against the target parameters.

#### 5.8.3. Metagenome assembly, binning, and quality control

To assess the practical ability to recover a pathogen’s genome, each of the 66 simulated metagenomes was subjected to a standard bioinformatics analysis pipeline as follows:

- Assembly: Raw simulated reads for each metagenome were assembled into contiguous sequences (contigs) using `metaSPAdes` (v4.2.0) (Nurk et al., 2017).
- Read alignment: To determine contig coverage, the simulated reads were mapped back to their respective assemblies using `BBMap`(`BBtools`v39.27) (Bushnell, 2014).
- Alignment processing: The resulting Sequence Alignment Map (SAM) files were sorted and converted to the BAM format using `Samtools` (v1.21) (Li et al., 2009).
- Binning: Contigs were binned into putative MAGs based on sequence composition and differential coverage using `MetaBAT2` (v2.17-66-ga512006) (Kang et al., 2019). The script `jgi_summarize_bam_contig_depths` (v2.17-66-ga512006) (Kang et al., 2019) was used to provide the required per-contig depth information to the binner.
- Quality control: The quality of the resulting pathogen MAGs was assessed using `CheckM2` (v1.1.0) (Chklovski et al., 2023), which calculates key metrics such as completeness and contamination. Taxonomic assignment of MAGs to confirm pathogen recovery was performed using the Genome Taxonomy Database Toolkit (`GTDB-Tk`, v2.4.1) (Chaumeil et al., 2022) with GTDB r220 (Parks et al., 2022).
- Throughout the downstream analysis of all 66 datasets, the total elapsed time and maximum resident RAM usage were monitored to evaluate the computational feasibility of the workflow.

### 5.9. Use of AI tools

The authors acknowledge the use of AI assistants for both code development and manuscript preparation. Algorithmic and documentation contributions (as detailed in the Methods section) were provided by Anthropic’s Claude-Sonnet 3.7. Structural feedback on the manuscript was additionally provided by Google’s Gemini 2.5 pro. All AI-assisted work was directed, reviewed, and validated by the human authors. The variable coverage simulation using overlapping sinewaves, realistic read length distribution functions for PacBio HiFi (log-normal) and ONT (mixed exponential-lognormal with heavy tail) platforms, and comprehensive code documentation were developed with AI assistance (Anthropic’s Claude-Sonnet 3.7). Algorithmic components were implemented by the AI based on biological specifications, while documentation was generated through analysis of the existing codebase. All AI-generated code and documentation were validated by the authors.

## 6. Acknowledgements

The work conducted by the U.S. Department of Energy Joint Genome Institute (https://ror.org/04xm1d337), a DOE Office of Science User Facility, is supported by the Office of Science of the U.S. Department of Energy operated under Contract No. DE-AC02-05CH11231.

## 7. Supplementary information

Complete timing and benchmarking data is presented in Supplemental Table S1. Protocol to install other simulators is presented Supplemental Information S1.

## 8. Conflict of interest

F.S. serves as CEO of SampleX. This work was not funded by, nor does it benefit SampleX. All other authors declare no competing interests

## 9. Software availability

`RandomReadsMG` available in the BBTools suite:

https://sourceforge.net/projects/bbmap/

For containerized deployment, a Docker image is also available from Docker Hub at

https://hub.docker.com/r/bryce911/bbtools.

BBTools is free for unlimited use under a LBNL open source license.

