## Supplemental Information for "Rapid terabase-scale simulation of realistic metagenomes for experimental design and pathogen detection with RandomReadsMG"

### Supplemental Information S1

#### 1. Protocol to install other simulators

```
conda create -n insilicoseq python=3.9
wget
https://repo.anaconda.com/miniconda/Miniconda3-latest-Linux-x86_64.sh
chmod +x Miniconda3-latest-Linux-x86_64.sh
./Miniconda3-latest-Linux-x86_64.sh
source ~/miniconda3/bin/activate
conda create -n insilicoseq python=3.9
conda activate insilicoseq
conda install -c bioconda insilicoseq
ls ../wgsim/wgsim/
realpath ../wgsim/wgsim/ref.fa
ls
ln -s ~/genomes/tid_2489212_1486976.Isoptericola.asm.fa ref.fa
time iss generate --genomes ref.fa --model hiseq --output r --n_reads
2000000 --cpus 32 --length 150

conda install -c conda-forge biopython
time iss generate --genomes ref.fa --model hiseq --output r --n_reads
2000000 --cpus 32 --length 150

conda install scipy
time iss generate --genomes ref.fa --model hiseq --output r --n_reads
2000000 --cpus 32 --length 150

conda install joblib
time iss generate --genomes ref.fa --model hiseq --output r --n_reads
2000000 --cpus 32 --length 150

conda install pysam
conda config --add channels defaults
```

```
conda config --add channels bioconda
conda config --add channels conda-forge
conda install pysam
time iss generate --genomes ref.fa --model hiseq --output r --n_reads
2000000 --cpus 32 --length 150

conda install requests
time iss generate --genomes ref.fa --model hiseq --output r --n_reads
2000000 --cpus 32 --length 150

conda install biopython=1.76
conda deactivate
conda create -n insilicoseq_py38 python=3.8
conda activate insilicoseq_py38
pip install InSilicoSeq
time iss generate --genomes ref.fa --model hiseq --output r --n_reads
2000000 --cpus 32 --length 150

iss -h
iss generate -h | grep length
time iss generate --genomes ref.fa --model hiseq --output r --n_reads
2000000 --cpus 32
reformat.sh in=r_R#.fastq
time iss generate --genomes ref.fa --model hiseq --output r --n_reads
4761904 --cpus 32
conda deactivate
```
